# Three distinct velocities of elongating RNA polymerase define exons and introns

**DOI:** 10.1101/044123

**Authors:** Qianqian Ye, Yoon Jung Kim, Hongyu Zhao, Tae Hoon Kim

## Abstract

Differential elongation rates of RNA polymerase II (RNAP) have been posited to be a critical determinant for pre-mRNA splicing. Molecular dissection of mechanisms coupling transcription elongation rate with splicing requires knowledge of instantaneous RNAP elongation velocity at exon and introns. However, only average RNAP elongation rates over large genomic distances can be inferred with current approaches, and local instantaneous velocities of the elongating RNA polymerase across endogenous genomic regions remain difficult to determine at sufficient resolution to enable detailed kinetic analysis of RNAP at exons. In order to overcome these challenges and to investigate kinetic features of RNAP elongation at genomic scale, we have employed global nuclear run-on sequencing (GRO-seq) method to infer changes in local RNAP elongation rates across the human genome, as changes in the residence time of RNAP. Using this approach, we have investigated functional coupling between the changes in local pattern of RNAP elongation rate at the exons and their general expression level, as inferred by sequencing of mRNAs (mRNA-seq). Our genomic level analyses reveal acceleration of RNAP at lowly expressed exons and confirm the previously reported deceleration of RNAP at highly expressed exons, suggesting variable local velocities of elongating RNAP that are potentially associated with different inclusion or exclusion rates of exons across the human genome.

**AUTHOR SUMMARY:** Understanding the mechanisms that enable high precision recognition and splicing of exons is fundamental to many aspects of human development and disease. Emerging data suggest that the speed of the elongating RNA polymerase affects pre-mRNA splicing; however, systematic genomic investigation of RNAP elongation speed and pre-mRNA have been lacking. Using a recently developed method for detecting synthesized nascent RNAs, we have inferred variable elongation rates of RNA polymerase II (RNAP) that are associated with included exons, introns and excluded exons, across the human genome. From this analysis, we have identified acceleration of RNAP at exons as a major determinant of exon exclusion across the genome, while confirming previous studies showing deceleration of RNAP at included exons.

## INTRODUCTION

The amazing diversity of the proteome of a human cell is governed by expression of a large number of transcript variants from a surprisingly small number of genes [1]. Annotation of functional units in the genome, exons and introns, would be an impossible task without the knowledge of cDNA sequences. Excluding the first and last exons, which exhibit additional sequence and biochemical features, short internal exons are largely indistinguishable amidst long introns whose average size that can easily exceed 100,000 nucleotides. Thus, how the spliceosome precisely and accurately finds the exons to splice is a great mystery. Understanding the molecular mechanisms that enable high precision recognition and splicing of exons is fundamental to many aspects of normal development, as well as pathologic perturbations that result in disease [2,3].

Multiple steps in pre-mRNA processing are coupled to transcription [4,5]. Alberto Kornblihtt and colleagues pioneered many of the earlier studies using minigene reporters have implicated kinetic coupling of transcription and splicing, in which slower transcription elongation rates modulate exon inclusion [6-9]. By swapping and testing different promoters driving a human fibronectin (FN) minigene reporter, the nature of the promoter, but not the strength of the promoter, was found to affect exon inclusion and exclusion, either by differential recruitment of splicing factors or by differential elongation rate of RNAP [6,7]. Analysis of the FN minigene model was further extended by use of a slow mutant of RNAP, which resulted in increased inclusion of the alternative exon. Most recently, the FN minigene reporter system was used to show exon inclusion can be modulated by RNAP C-terminal domain (CTD) phosphorylation upon UV irradiation and CTD phosphorylation can be tentatively associated with changes in RNAP elongation rate [8]. Although these studies using the artificial FN minigene system have implicated kinetic coupling of transcription and splicing in which variable elongation rate modulate exon inclusion, it remains unclear whether such mechanism plays a critical role in expression of endogenous genes within their proper chromatin environment, and if so, how widespread this kinetic coupling mechanism is across the human genome.

Recent advances in functional genomics and bioinformatics approaches have enabled molecular insights that are sometimes unattainable from analysis of individual genes and loci by aggregate analyses of many similar regions across the genome to reveal global patterns and trends. An average *internal* exon, excluding the first and last exons, is 147 nucleotides long, an exact length of DNA found around a nucleosome. The length of an exon being the exact length of a nucleosomal DNA [10] does not seem coincidental. Recent genome-wide studies demonstrate nucleosomes as a fundamental organizing unit and substrate of genetic processes. Exons may have evolved to just one nucleosome’s length to facilitate their identification and splicing [11]. Chromatin immunoprecipitation (ChIP) coupled to high throughput sequencing (ChIP-seq) analyses of histones have revealed increased nucleosomal density at exons, compared to surrounding intronic sequences [12-14], suggesting epigenetic coding of exons by differential positioning of nucleosomes. In an earlier chromatin immunoprecipitation coupled to DNA microarray (ChIP-chip) analysis of RNAP, Pam Silver and colleagues described an increase in RNAP density at exons genome-wide [15]. This increased occupancy of RNAP at exons has been interpreted as decreased rate of RNAP elongation at the exons. Potentially, this local pausing of RNAP at exons is due to the positioned nucleosomes at the exons that serve as “speed bumps” [16]. Consistent with this observation, nucleosomes have been shown to act as physical barriers to RNAP elongation in single molecule studies *in vitro* [17].

After normalizing for the observed increased nucleosomal density, significantly elevated levels of several histone modifications are still detected at exons across the genome, suggesting that positioned nucleosomes at exons are preferred substrates for histone modifying enzymes. DNA methylation is also selectively enriched at the exons [18], suggesting that nucleosomes are the relevant *in vivo* substrates for the DNA methyltransferases [19]. Among the histone modifications analyzed thus far, trimethylation of lysine residue 36 of histone H3 (H3K36me3) exhibits the strongest correlation with the internal exons [12,13,20], suggesting that epigenetic modifications of histones and DNA may facilitate recognition of exons by slowing down the elongating RNAP. However, a systematic investigation of RNAP elongation rate at exons and introns across the human genome has been limited.

To investigate elongation velocity of RNAP across the human genome at a resolution that enables systematic analysis at exons, introns and other features, we have used global nuclear run-on sequencing (GRO-seq) method to infer “residence time” of RNAP across the genome and local changes in RNAP elongation rate. Combining this approach and deep sequencing of mRNAs, we report that variable RNAP elongation rates are associated with excluded exons, introns and included exons. We further investigate this global pattern of RNAP elongation rate with pre-mRNA splicing to implicate splicing and transcription factors in modulating the elongation velocity of RNAP.

## RESULTS

To infer RNAP elongation rates across the genome, we have performed GRO-seq [21], in a trio of normal diploid human cell lines (H1 embryonic stem cells, IMR90 fibroblasts and MCF10A breast epithelial cells). This method allows the detection of nascent transcripts from the engaged RNAP, and thus can be used as a measure of elongating RNAP density across the genome, averaged over millions of nuclei. In general, GRO-seq tag density is inversely correlated with the RNAP elongation rate [21] and is reflective of the “residence time” of RNAP at a particular genomic region. We aligned the resulting GRO-seq reads, as well as publically available H3K36me3 ChIP-seq reads relative to Ensembl annotation and averaged the mapped tag density of all internal exons and flanking sequences. GRO-seq tag density was enriched at exons and centered around the exon midpoint (Figure 1A-C). H3K36me3 tag density was also enriched in exons and peaked slightly downstream of the exon midpoint in all three cell lines (Figure 1A-C). These results are consistent with the previous studies [12,15,22,23] that show exonic pausing of RNAP [24] and epigenetic definition of exons [20]. The slight spatial difference between GRO-seq peak and H3K36me3 peak positions relative to the exon midpoint is consistent with the notion that nucleosome limits RNAP elongation [17].

**Figure 1.**
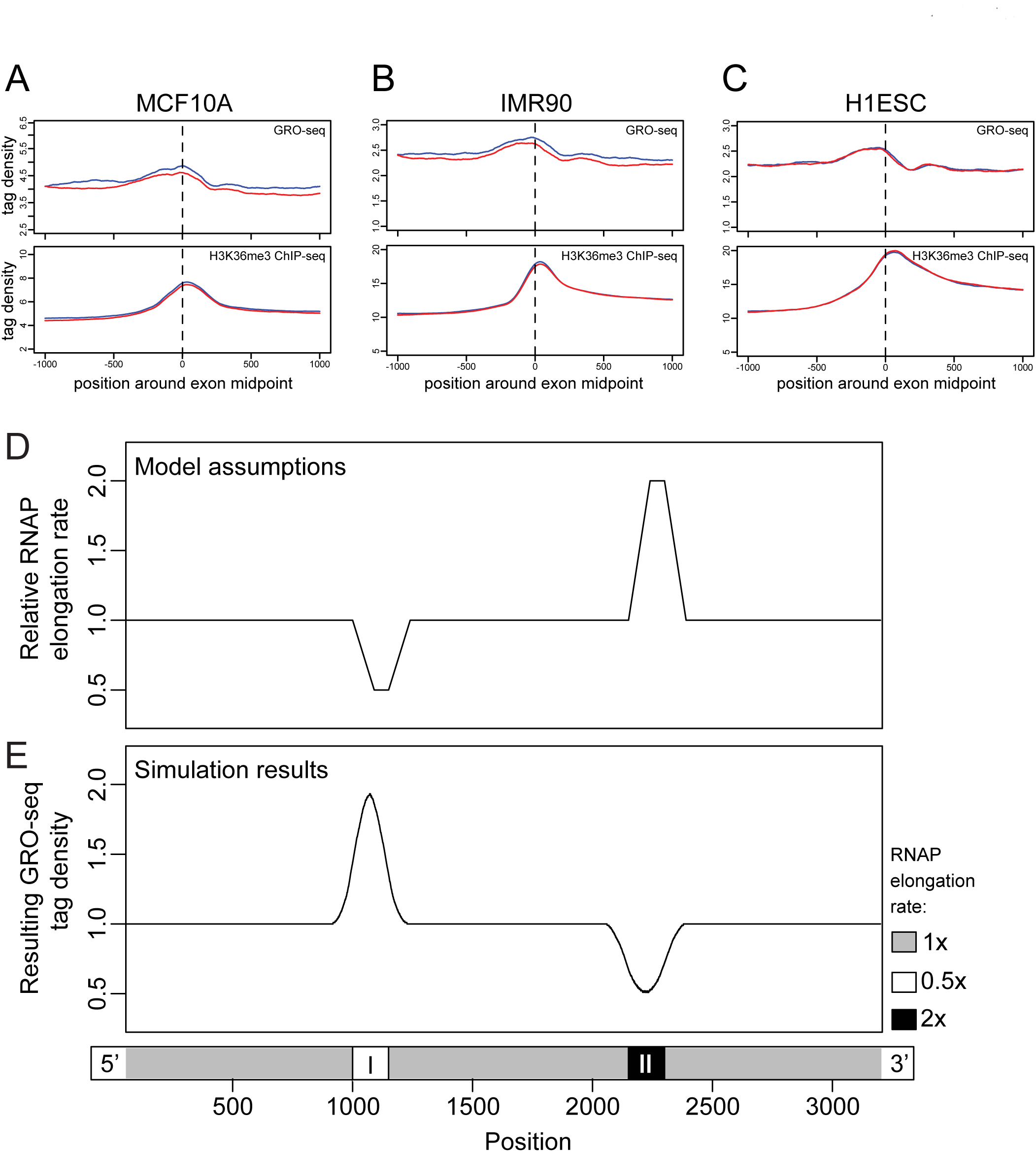
Patterns of GRO-seq tag density are related with changes of RNAP elongation rate. (A-C) Average mapped sequence tag densitites in MCF10A (A), IMR90 (B) and H1ESC (C) are shown. GRO-seq tag density and H3K36me3 ChIP-seq tag density of internal exons were averaged around exon midpoints (vertical dashed line) in the three cell lines, respectively. The red curves represent exons on the positive strand, and the blue curves represent exons on the negative strand. Average internal exon lengths are 146.7 bp in MCF10A, 146.7 bp in IMR90, and 146.2 bp in H1ESC. (D) Profile of RNAP elongation rate as the model assumption. Three 1 kb DNA segments are separated by two 150 bp regions, I and II, with vertical dashed lines showing their start sites, midpoints and end sites from left to right. RNAP elongation rate changes in region I and region II to achieve two-fold deceleration and acceleration, respectively, and changes back graduately at the downstream of the two regions. (E) Simulated GRO-seq tag density level based on the profile of RNAP elongation rate showing a peak in region I where elongation rate is doubled and a dip in region II where it is halved. The peak and the dip are symmetric with respect to the midpoint of I and II, respectively.

### Computational modeling of nascent RNA mapping

To interpret the spatial difference in GRO-seq data in greater detail and to investigate if GRO-seq data can infer local kinetic properties of the elongating RNAP, we have developed an *in silico* model to predict GRO-seq tag density along an artificial segment of DNA. We incorporated the experimentally determined parameters of the GRO-seq protocol [21] and defined arbitrary changes in the RNAP elongation rate along a transcribed segment to model GRO-seq tag densities that would result from variable RNAP elongation rates. Our model DNA contained three large 1,000 bp segments separated by two small internal 150 bp segments. We have defined relative RNAP elongation rates on the three 1000 bp segments as “1”, and on the two 150 bp segments as “0.5” and “2”, respectively (Figure 1D). We also assumed constant RNAP loading and promoter escape rates at the 5’ end. 10,000 entities were simultaneously simulated and the relative GRO-seq tag density was calculated at single nucleotide resolution (see Methods for details). An approximate two-fold enrichment of GRO-seq tags was obtained at the region with two-fold decrease of RNAP elongation rate (Figure 1E), reflecting the increased RNAP residence time. An opposite pattern of GRO-seq tag density at the region where RNAP moves twice as fast was determined from our simulation (Figure 1E), reflecting the decreased RNAP residence time. These simulation results indicate that GRO-seq tag density is inversely correlated with RNAP elongation rate, reflecting residence time of RNAP at the region, and can potentially be used to infer relative changes in local RNAP elongation rate. The shift of the simulated GRO-seq tag density peak from the peak of underlying RNAP elongation rate agrees well with our experimental results (Figure 1A-C) and points to functional association between H3K36me3 and RNAP transcription elongation.

### Identification of distinct patterns of RNAP elongation velocity using GRO-seq data

In order to analyze the fine-scale patterns within our GRO-seq density data at exons of transcribed genes, we divided the pool of internal exons into ten groups by exonic GRO-seq tag density mapped on either Watson or Crick strands of the genome (blue or red lines, respectively, in Figure 2). Group 1 contains exons with exonic GRO-seq tag density equal 0, and the rest of the exons were equally divided into 9 groups by exonic GRO-seq tag density. Averaged GRO-seq tag densities at single nucleotide resolution of each group show modest difference in intronic GRO-seq tag density level among the groups, which range from 1 to 13 (Figure 2). However, this stratification of our GRO-seq data revealed much more dynamic changes in GRO-seq tag density at the exons relative to introns. Three obvious patterns of exonic GRO-seq tag density relative to the intronic GRO-seq tag density can be classified that match our *in silico* model. First pattern (dip) is marked by a decrease in GRO-seq tag density at the exon from the flanking introns (Figure 2, group 1 to 5, upper panel), suggesting that RNAP is accelerating through these exons. Second pattern (flat) shows the same tag density as the bordering intronic level (Figure 2, groups 6 and 7, upper panel). Third pattern (peak) is characterized by an increase in GRO-seq tag density from the intronic level (Fig 2, group 8 to 10, upper panel), suggesting that RNAP is decelerating at these exons. Similar GRO-seq pattern was observed in all the three cell lines we have examined (Figures S1, S2 and S3). In addition, we confirmed that the difference in GRO-seq pattern was independent of GC content (Figure S4), Br-UTP labeling efficiency in GRO-seq experiments (Figure S5) and mappability of the 35-nucleotide sequence tags (Figure S6).

**Figure 2.**
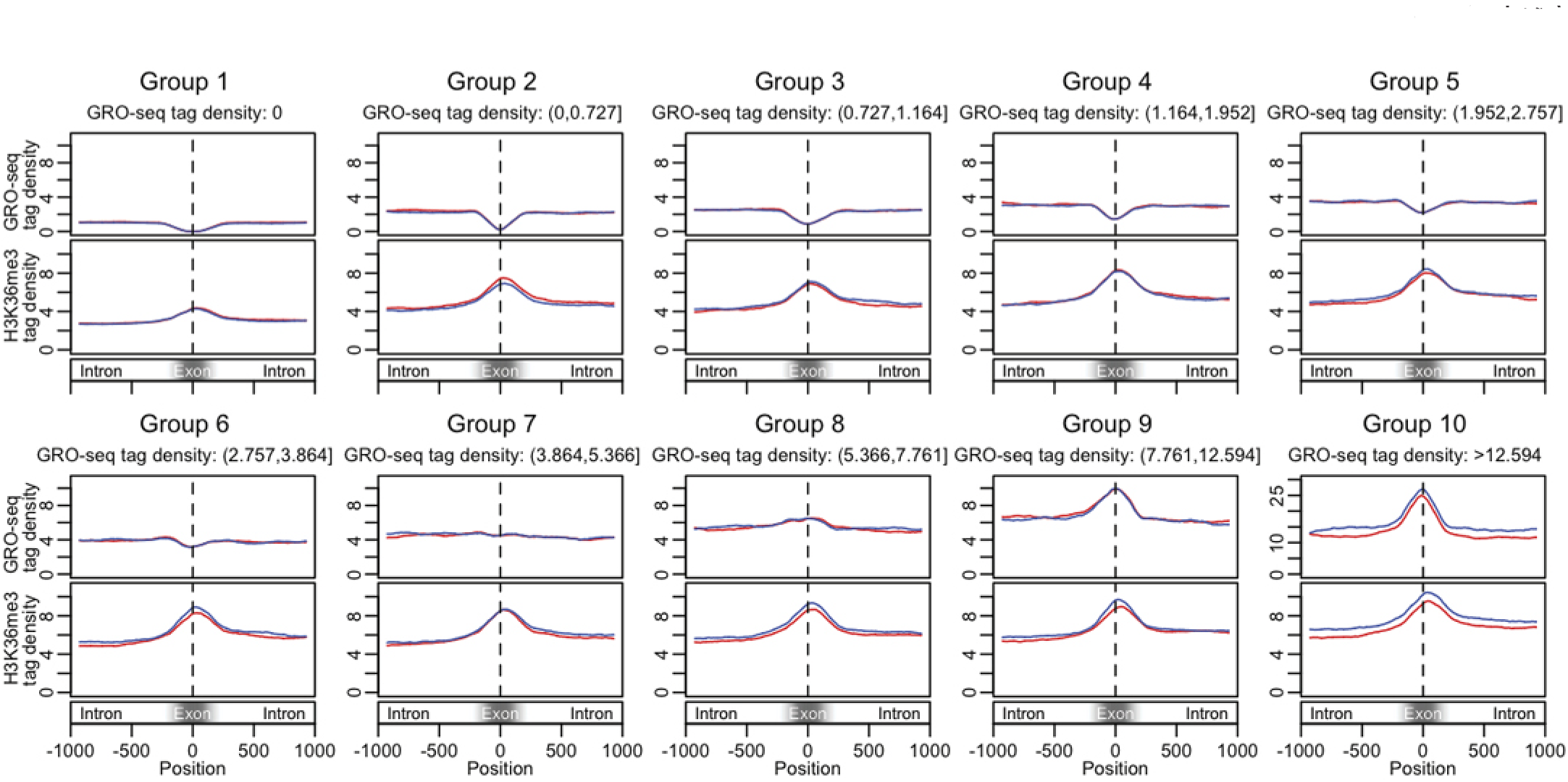
GRO-seq tag density reveals three distinct patterns at exons in MCF10A. GRO-seq reads aligned around midpoints of exons indicated by black vertical dashed lines (upper panel of each group) and averaged H3K36me3 ChIP-seq tag density in the same group of exons (lower panel of each group). Exonic GRO-seq tag density range of each group is indicated below the group names. Red curves and blue curves correspond to exons on positive strand and negative strand, respectively. Patterns of GRO-seq tag density changes continuously and gradually from one group to another. Three distinct patterns (dip, flat behavior and peak) of GRO-seq tag density are observed within exons in groups with low (group 1 to 5), medium (group 6 and 7) and high (group 8 to 10) exonic GRO-seq tag densities, while intronic GRO-seq tag densities are about the same (except group 1 and 10). The patterns are symmetric to the midpoints. On the other hand, H3K36me3 peaks locating in exons are observed in all 10 groups, with difference in relative peak height, but not in general pattern. In comparison with the symmetry of GRO-seq peaks, the H3K36me3 peaks shift slightly towards to 3’ direction of exon midpoints. Also see Figure S2 for averaged tag densities around 3’ end and 5’ end of exons.

### Development of *GRO-seq pattern score* to quantify spatial differences in RNAP elongation velocity

To capture and analyze the spatial information of our GRO-seq data, we have developed a simple calculation which we defined as GRO-seq “pattern score” for each exon as a ratio of exonic tag density to tag density of flanking introns (Figure 3A). The GRO-seq pattern score also serves to normalize GRO-seq tag density at the exons to the densities found that flanking introns. Distributions of GRO-seq pattern scores were consistent with averaged GRO-seq tag density patterns (Figure S7A). In groups with lower exonic GRO-seq tag densities, more exons have GRO-seq pattern scores lower than 1 (Figure S7A, group 1 to 5), corresponding to decreased GRO-seq level in exonic regions compared with the flanking introns; and in groups with higher exonic GRO-seq tag densities, more exons have GRO-seq patterns scores higher than 1 (Figure S7, group 8 to 10), corresponding to enriched GRO-seq tags in exonic regions. This observation is also consistent among all three cell lines examined (Figure S7B and S7C). Although we define three major categories using GRO-seq pattern score normalized to the flanking intronic GRO-seq tag densities, GRO-seq pattern scores show a continuous distribution (Figure S8)

**Figure 3.**
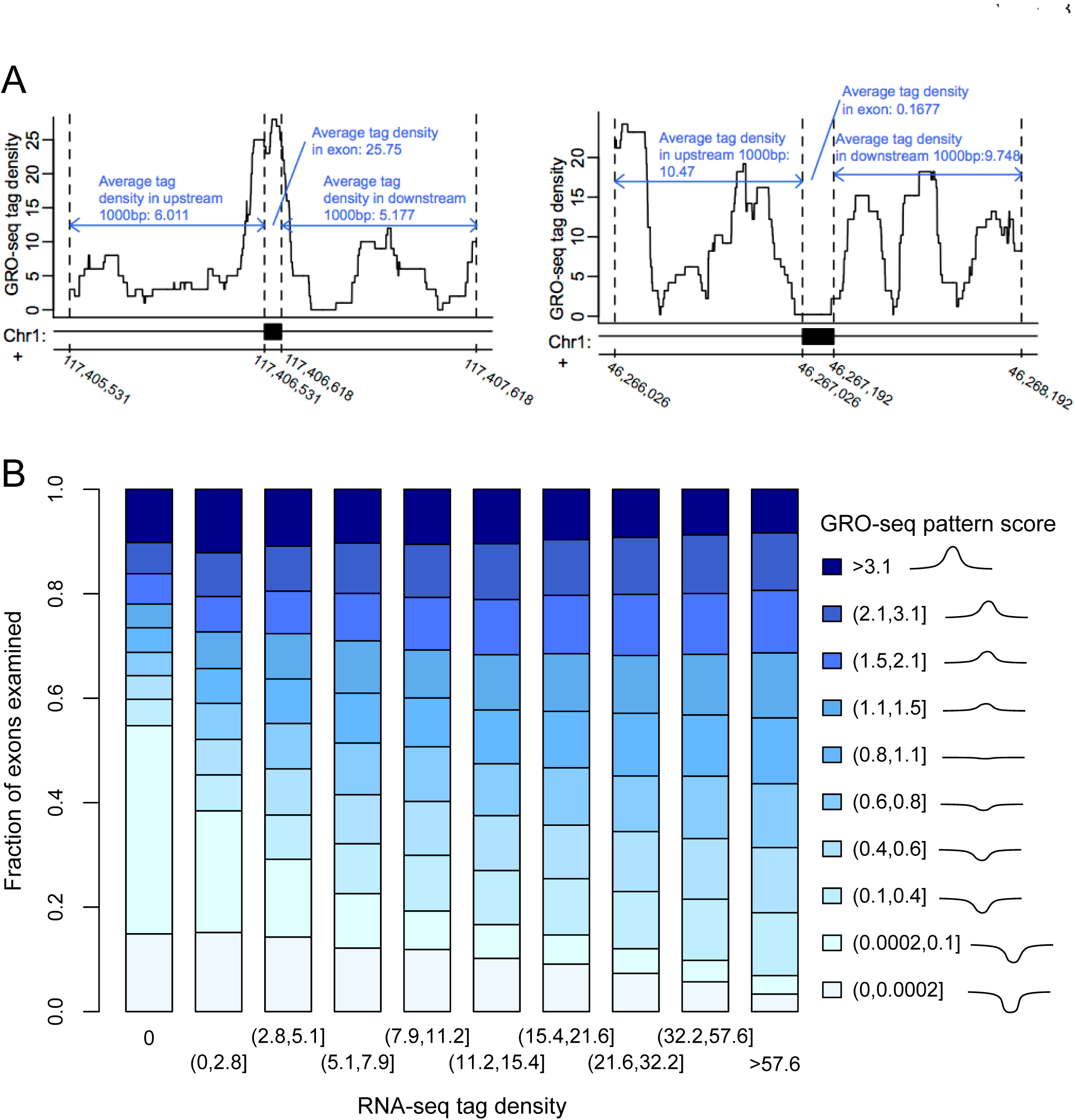
GRO-seq tag density pattern correlates with mRNA level in MCF10A. (A) Two examples of calculating pattern score of GRO-seq in MCF10A. GRO-seq pattern score of the exon in left panel is 25.75*2/(6.011+5.177)=4.603, and the one in right panel is 0.1677*2/(10.47+9.748)=0.01659. (B) Bar plot shows distributions of GRO-seq pattern scores in 10 groups divided by exonic mRNA-seq tag density. Each group of mRNA-seq tag density is represented by one bar. Exons were also classified into 10 groups of continuous value ranges of GRO-seq pattern score as mentioned in the main text, indicated by the legend on the right of the plot. The proportions of exons within different GRO-seq pattern score value ranges show a shift among 10 groups of mRNA-seq tag density.

### mRNA-seq analysis to determine the fate of exons with different GRO-seq pattern scores

In order to characterize the exonic RNAP elongation rates inferred from GRO-seq pattern score and exon expression level, we performed mRNA sequencing (mRNA-seq) analysis to measure relative representation exons in polyadenylated RNAs [25,26]. We determined the relative expression level for each exon, defined as exonic mRNA-seq tag density that perfectly correlated with the standard metric of RNA abundance, RPKM (RNA sequence tags per kilobase) [26]. GRO-seq pattern score correlated with exon expression level and mRNA level in all three cell lines (Figure 3B and Figures S9 and S10). Specifically, in MCF10A, about 70% of exons with zero mRNA-seq tags recovered have dips of GRO-seq pattern in the exonic regions (Figure 3B, left most bar). This percentage drops gradually as the observed mRNA level increases (Figure 3B, right most bar). The correlation between GRO-seq tag density pattern and mRNA level (Figure S11) suggests that exon inclusion may be regulated by elongation rate, i.e. slower elongation rate in exons than in flanking introns correlates with exon inclusion, while faster exonic elongation rate correlates with exon exclusion [6,7,22]. In addition, we did not observe similar mRNA level-related shift in RNAP ChIP-seq pattern score (Figures S12, S13), suggesting that GRO-seq may be more sensitive for characterizing engaged RNAP position compared with conventional ChIP-seq [27,28].

Based on this observation of distinct GRO-seq pattern at exons relative to introns reflecting changes in RNAP elongation rate, we examined GRO-seq tag densities of two sets of exons arising from the same gene: one set with zero mapped mRNA-seq tags from the same annotated genes (excluded exons), and the other set with more than zero mapped mRNA-seq tags (included exons). We found that difference between the GRO-seq at individual exons from the same gene showed that included exons tend to have higher GRO-seq tag densities than the excluded exons of the same annotated gene (Figure S14A). When we plot distribution of difference in GRO-seq tags between included and excluded exons, we also observed that the included exons exhibit higher GRO-seq tag densities than the excluded exons from the same gene (Figure S14B). We have investigated whether we have enough resolution to determine if individual exons can be confidently assigned its mRNA inclusion rates as well as reliable GRO-seq pattern score. At the current sequencing depth and read length, determination of relative exclusion rate of individual exons in the expressed mRNA remains difficult, as well as ascertainment of GRO-seq patterns at individual exons. This limited resolution at individual exons arises from the nature of short read length of mRNA-seq precluding meaningful and accurate measures of exon inclusion/exclusion rate for most of the exons and relatively low-depth coverage of GRO-seq at current throughput used in our analysis. With increases in sequencing depth and length, it may be possible to make direct comparison of exon inclusion/exclusion rates with the GRO-seq pattern score. Thus, our observation at the genomic, aggregate level should be taken with some caveats concerning exon inclusion and exclusion rates.

### Comparison of GRO-seq with ChIP-seq data

One of the key histone modifications associated with transcription elongation and with exons is H3K36me3 [13,29]. We investigated how patterns of GRO-seq tag density is correlated with the patterns of H3K36me3 at exons. In all three cell lines, H3K36me3 levels corresponded to the larger GRO-seq pattern scores for exons within the Group 9 and 10 (Figure S2). In other groups of exons, however, we did not detect correspondence between GRO-seq and H3K36m3 patterns (Figure 2 and Figure S14). This result was further confirmed by analyzing H3K36me3 pattern score distributions over the 10 groups with different exonic GRO-seq tag density ranges, which correspond to different GRO-seq patterns as we have already described in Figure 2. Unlike the GRO-seq data, distribution of H3K36me3 pattern score did not show a similar shift as the one of GRO-seq pattern score (Figures S8A and S15). In addition, we did not observe a similar correlation between mRNA level and H3K36me3 pattern (Figure 4 and Figure S16). Lastly, we did not observe strong association between GRO-seq pattern and H3K36me3 pattern or other epigenetic patterns (H4K20me3, H3K9me3 and DNA methylation) (data not shown). Combined, these results suggest that nucleosome and its modifications at exons are not be sufficient to determine GRO-seq patterns and transcription elongation rate at exons.

**Figure 4.**
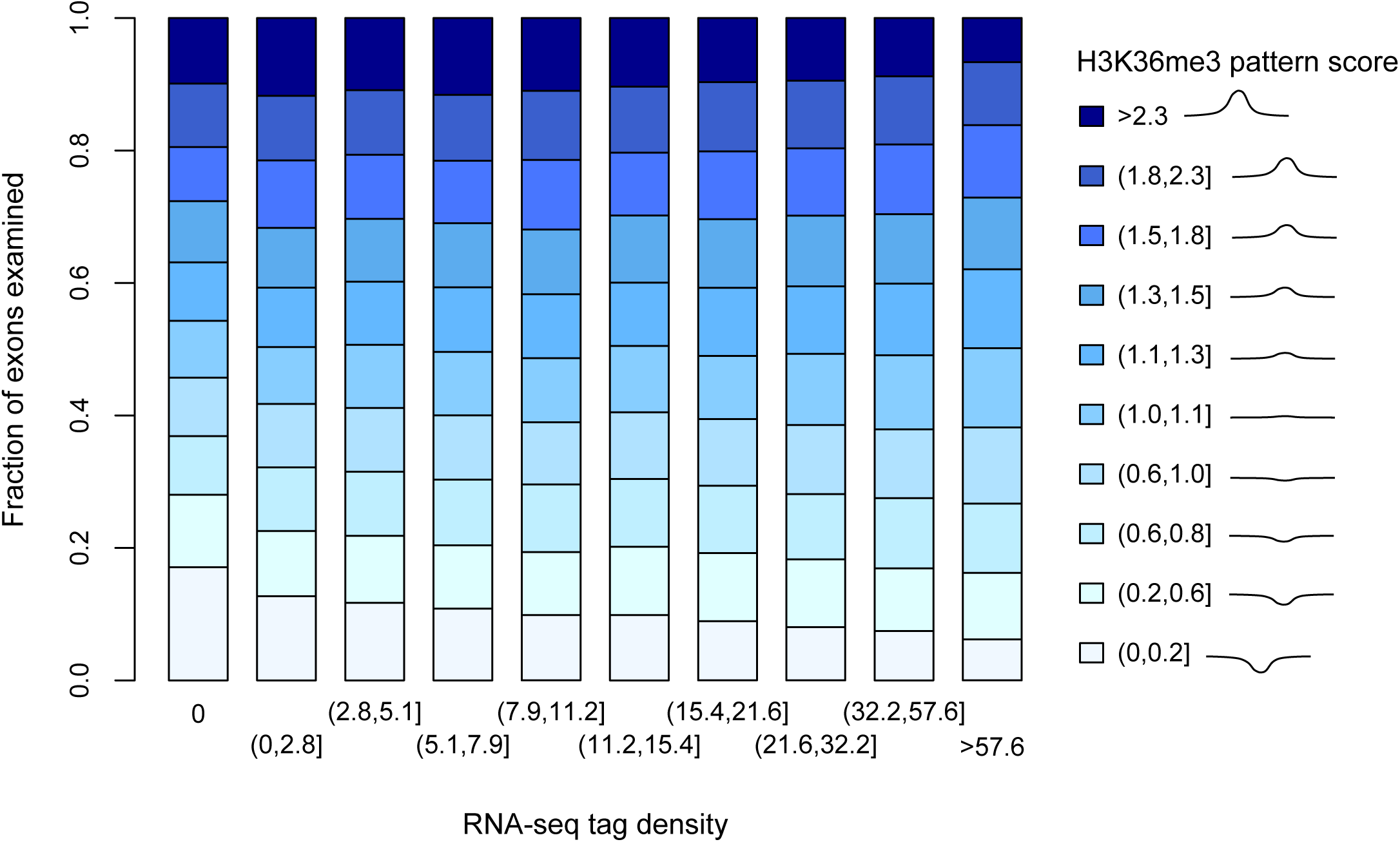
H3K36me3 pattern scores are not related with mRNA level in MCF10A. Bar plot shows distributions of H3K36me3 ChIP-seq pattern scores in 10 groups divided by exonic mRNA-seq tag density. Each group of mRNA-seq tag density is represented by one bar. Exons were also classified into 10 equal-sized groups of continuous value ranges of H3K36me3 ChIP-seq pattern score, as indicated by the legend on the right of the plot. The proportions of exons within different H3K36me3 ChIP-seq pattern score value ranges show a shift among 10 groups of mRNA-seq tag density.

### Statistical modeling of RNAP elongation velocity and gene expression

Based on the patterns of GRO-seq and ChIP-seq data, we explored importance of RNAP elongation rate as inferred by GRO-seq and histone and DNA methylation in determining exon inclusion inferred from mRNA-seq data. We used variable selection of a random forest model to explore relative contribution of GRO-seq and epigenetic marks to exon inlcusion, which was estimated based on the exon abundance from mRNA-seq data (Table 1). A random forest model was built to predict exon inclusion and contained exonic intensity and pattern score of six epigenetic features that correlate with exon definition, and they are GRO-seq, H3K36me3 [12-14], H4K20me3 [12-14], H3K9me3 [30], DNA methylation [18,19], RNAP positioning [12,15,22,23] and exon length as a negative control. Backward variable selection was applied to remove variables with lower importance than exon length in predicting exon inclusion, resulting in 6 variables remained within the reduced model (Table 1). Compared with the prediction accuracy of the full model, which was 0.8199±0.0035 (Cohen's kappa coefficient: 0.4205±0.0096), the reduced model reached prediction accuracy 0.8138±0.0033 (Cohen's kappa coefficient: 0.4073±0.0092), showing only minor loss of prediction power. Among the remaining six variables, GRO-seq exonic tag density showed the highest importance, followed by GRO-seq pattern score, while H3K36me3, DNA methylation and H4K20me1 were less important for the model. These results show some consistency with recent study showing relationship between mRNA splicing and H3K36me3 levels [31]. Furthermore, our results suggest that RNAP elongation rate (as determined by GRO-seq tag density and pattern score) has a more direct effect on exon inclusion and may incorporate histone and DNA methylation information at the exons to refine mRNA splicing.

**Table 1.** Variable importances measured in mean decrease of accuracy in the random forest model of exon inclusion in IMR90. The first column lists variables tested in the model. The second and third columns list the mean decrease in accuracy of the model upon removal of the variable, employing the entire set of variables (Full Model) or a reduced set (Reduced Model).

| Variable | Importance (Mean Decrease of Accuracy) |  |
| --- | --- | --- |
|  | Full Model | Reduced Model |
| GRO-seq exonic tag density | $0.0588 \pm 0.0011$ | $0.0750 \pm 0.0014$ |
| H3K36me3 exonic tag density | $0.0276 \pm 0.0006$ | $0.0236 \pm 0.0011$ |
| GRO-seq pattern score | $0.0204 \pm 0.0006$ | $0.0267 \pm 0.0006$ |
| DNA methylation level | $0.0112 \pm 0.0004$ | $0.0166 \pm 0.0008$ |
| DNA methylation pattern score | $0.0103 \pm 0.0004$ | $0.0156 \pm 0.0008$ |
| H4K20me1 exonic tag density | $0.0101 \pm 0.0006$ | $0.0096 \pm 0.0006$ |
| H3K36me3 pattern score | $0.0095 \pm 0.0003$ | |
| exon length | $0.0072 \pm 0.0003$ | |
| H3K9me3 exonic tag density | $0.0054 \pm 0.0003$ | |
| RNAPII exonic tag density | $0.0053 \pm 0.0003$ | |
| RNAPII pattern score | $0.0050 \pm 0.0002$ | |
| H4K20me1 pattern score | $0.0047 \pm 0.0001$ | |
| H3K9me3 pattern score | $0.0039 \pm 0.0002$ | |

### Accelerators and decelerators of RNAP elongation

Sequence features may also contribute to changes in RNAP elongation velocity at the exons. We asked whether there are differences in splicing factor binding motifs at or near exons with distinct GRO-seq pattern scores within each cell line. We analyzed the densities of putative known splicing factors binding sites annotated in the SFmap [32,33] for the exons of expressed transcripts in each of the three cell lines. Putative motifs of 17 known splicing factors [33] were analyzed. One-sided Wilcoxon rank sum tests were performed to test for difference in the mapped motifs at exons that exhibit distinct GRO-seq pattern scores corresponding to acceleration (pattern score ≤ 0.5) or deceleration (pattern score >2) of RNAP elongation rate in each of the three cell lines. Results from both exonic regions (Figure 5A) and flanking intronic regions (Figure 5B) revealed several splicing factors that exhibit differences in motif density in three cell lines. We found that splicing factor binding site motifs are more abundant at or near exons with faster RNAP elongation rate, suggesting that these splicing factor binding site motifs may refine the epigenetic definition of exons to facilitate exclusion of nucleosomally marked exons. In addition, H1 ESC displayed more contrasting densities of splicing factor binding site motifs than IMR90 and MCF10A cells, suggesting that embryonic stem cells might have a distinct splicing regulatory network from the two somatic cells as previously suggested [34].

**Figure 5.**
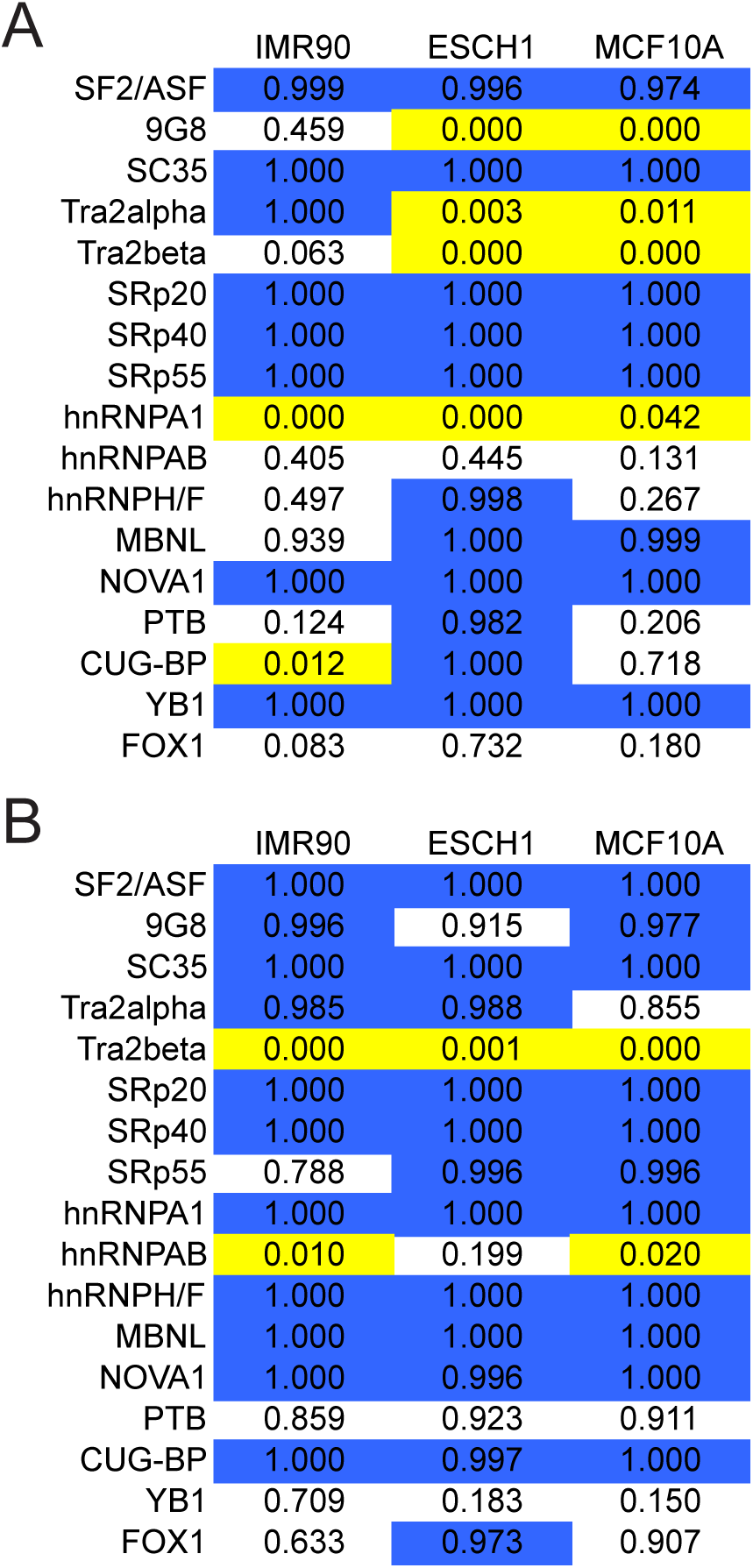
Differential enrichments of putative known splicing factor motifs in exons with cell-type-specific GRO-seq patterns. P values of one-sided Wilcoxon rank sum tests for length-normalized frequencies, i.e. densities, of exonic (A) or intronic (B) splicing factor motifs located no further than 100 bp from splicing sites. Wilcoxon rank sum tests were performed between exonic and intronic motif densities of exons with exonic GRO-seq pattern scores > 2 and those with exonic GRO-seq pattern scores ≤ 0.5 in each cell line. The p value of each test is shown in the palette. Significant p values were further defined by Benjamini and Hochberg FDR procedure[48] at the 0.05 level and indicated by colored background: yellow indicates motifs are enriched at exons with GRO-seq pattern scores > 2 (decelerators), and blue indicates motifs are enriched exons with GRO-seq pattern scores ≤ 0.5 (accelerators).

We extended this splicing motif enrichment analysis to exons that display different RNAP velocities between cell lines, to identify potential cell type specific regulators that may contribute to different RNAP velocities at given exons. We analyzed exons that show distinct GRO-seq pattern scores in pairwise comparisons of H1 ESC, IMR90 and MCF10A (Figure S17). Results from both exonic regions and flanking intronic regions revealed a number of splicing factor binding sites that exhibit differences in density among the cell types (Figure S18). In particular, difference of expression level (as determined by mRNA-seq) of Tra2beta gene among the three cell lines was consistent with the difference in their exonic motif density among the cell lines examined (Figure S18 and Table S2). Similarly, hnRNPA1 showed consistent expression patterns and intronic motif density (Figure S18 and Table S2) across the three cell lines. Again, differences in splicing factor motif density between ESC H1 and the either of the two cell lines were more pronounced than between the two differentiated somatic cell lines, IMR90 and MCF10A.

We performed a more systematic analysis of sequence motif densities by exhaustive enumeration of 6, 7 and 8-mers at exons with different RNAP elongation rates among the three cell types. We identified additional sequence elements that associate with different RNAP elongation rate at exons. The resulting significant n-mers of potential RNAP accelerators/decelerators from the search were compared against databases of known *cis*-regulatory motifs (TRANSFAC [35] and JASPAR [36]) using STAMP [37] (Tables S3, S4, S5 and S6). In light of recent work implicating transcription factor binding sites in pausing of RNAP and induction of exon inclusion [38], our results suggests a number of diverse transcription factor binding sites may play a role in regulating RNAP elongation rate at exons. Thus, these enriched sequence features at distinct GRO-seq patterns suggest that splicing and transcription factors contribute to the regulation of transcription elongation rate and to the coupling of transcription elongation with mRNA splicing [31,39].

## DISCUSSION

Previous efforts in measuring the elongation rate of RNAP across endogenous genes employed a range of methods including reverse transcription PCR, nuclear run-on assays and fluorescence in situ hybridization. The most recent method developed by Singh and Padgett [40] uses DRB to transiently stall the RNAP at the promoter, then release the stalled RNAP by washing away the drug and collecting the RNA at various time points and performing real-time RT-PCR quantification of the nascent RNA. Using this method, RNAP elongation rate was estimated at about 3.8 Kb per minute. However, this is an average speed of RNAP across a large genomic region and several domains. Based on this average rate, RNAP will transit through an exon in about 2.3 seconds, suggesting that this standard approach for investigating RNAP transit times would require high temporal and spatial resolution to detect potential acceleration or deceleration that may occur at exons. Due to these challenges in measuring variations of local RNAP elongation rate, our understanding of genome-wide relationship between variable RNAP elongation rates and pre-mRNA splicing has remained obscure.

Here, we described an approach using global run-on sequencing (GRO-seq) methods for analyzing changes in local rates of RNAP elongation across the human genome. We have applied this method to human embryonic stem cells, differentiated lung fibroblasts and breast epithelial cells. By combining GRO-seq with mRNA-seq experiments, we have begun to investigate pre-mRNA splicing and RNAP elongation velocity. Our results suggest highly variable RNAP elongation rates that are associated with different expression levels of exons as inferred by mRNA-seq. We also find that transcription elongation associated modification, H3K36me3, is invariantly marked at exons regardless of the changes in RNAP elongation rate at exons, suggesting that modulation of RNAP elongation rate and splicing may function independently of epigenetic definition of exons at global level. Consistent with this view, our computational analyses identified distinct sequence features that are associated with acceleration and deceleration of RNAP. Known splicing factor motifs as well as transcription factor motifs are differentially enriched at exons with distinct RNAP elongation rates inferred by GRO-seq patterns at exons. Functional studies are in progress to investigate and explore sequence features associated with acceleration and deceleration of elongating RNAP.

Our study provides a potentially simple analytic framework for a systematic and genome-wide analysis of RNAP elongation rate as it relates to pre-mRNA splicing. By leveraging this analytical framework, we hope to extend the current work to analyze mechanisms that couple pre-mRNA splicing to transcription elongation. A major limitation of our current approach is the lack of accurate genome-wide measurements of exon inclusion/exclusion rates, which precludes our analysis from exploring various mechanisms that can modulate coupling of RNAP elongation rate to pre-mRNA splicing at individual exons, but emerging new sequencing technologies may help us to overcome this problem [41]. Nonetheless, at a broad genomic level, our study suggests that kinetic control of RNAP elongation [42] may be a general mechanism for regulating pre-mRNA splicing across the human genome and provides a new analytic framework for deciphering the mechanisms of kinetic coupling of transcription and pre-mRNA splicing.

## METHODS

### Cell culture

MCF10A cells were cultured with DMEM/F12 media supplemented with 2.5% Horse serum, 0.5ug/mL hydrocortisone, 10ug/mL Insulin, 20ng/mL EGF and 100ng/mL cholera toxin. IMR90 cells were cultured in MEM with Earles salts with 10% FBS. H1 embryonic stem cells (WiCell) were cultured on feeder-free matrigel coated plates in Dulbecco's modified Eagle medium (DMEM)/F12, supplemented with 1% MEM-nonessential amino acids (Invitrogen), 1 mM L-glutamine, 1% penicillin-streptomycin, 50 ng/mL basic fibroblast growth factor (Millipore), N2 supplements (1X), and B27 supplements (1X) (Invitrogen).

### Sequencing Libraries

Global run-on and library preparation for sequencing was performed based on the previously published method [21]. Briefly, intact nuclei were isolated and incubated for 5 minutes at room temperature with nucleosides and bromo-UTP analog to allow RNA polymerase to incorporate and nucleosides in a nuclear run-on reaction. The nascent RNA from the run-on reaction was isolated and subjected to immunoaffinity purification using the Br-dUTP antibody (Santa Cruz Biotech). After first round of purification of nascent RNA, the resulting RNA was ligated with 5' small RNA adaptor RNA (Illumina) and subjected to second Br-UTP purification. The resulting RNA is further ligated using 3' adaptor RNA and subjected to third round of purification. The final purified RNA was reverse transcribed and PCR amplified and size selected (150-250bp) using polyacrylamide gel electrophoresis to generate the final sequencing libraries. We performed the short RNA sequencing protocol of the GRO-seq libraries per instructions from the manufacturer (Illumina).

For ChIP-seq assays, we have followed the ChIP protocol previously published [43]. The ChIP DNA was further processed by the Yale Center for Genomic Analysis following the instructions from Illumina. For mRNA-seq, total RNA was processed by the Yale Center for Genomic Analysis for Whole Transcriptome Analysis (mRNA-seq) following the instructions from the manufacturer (Illumina).

### Simulation of GRO-seq

Because we were interested in internal exons, architecture of the gene segment used for simulation contains three 1 kb segments separated by two 150 bp regions, named region I and region II in the 5’-to-3’ direction. The base positions were named from 1 to 3,300 in the 5’-to-3’ direction. We assumed that starting from the 5’ end, transcription elongation rate *v*_0_ is a constant before it hits region I. In region I, the rate decreases with constant deceleration *-α*_*I*_ until it equals ½*V*_0_. From the 3’ end of region I, the rate starts to increase with constant acceleration *α*_*I*_ back to *v*_0_. The constant was derived by satisfying the symmetry of GRO-seq tag density pattern around midpoint of region I. The elongation rate around region II was derived similarly, with the highest rate in region II being 2*ν*_0_. By assuming constant RNAP loading rate at 5’ end, i.e. constant temporal distance (*T*) between each two adjacent RNAP molecules, RNAP positions along the gene segment is derived from transcription elongation rate and the position of the RNAP molecule most close to 5’ end of the gene segment (*P*_*5*_). 10,000 identical gene segment entities were simulated. RNAP positions of each entity were calculated with the assumption of uniform distribution of *P*_*5’*_ in [1, *Tv*_0_]. GRO-seq tags were then generated as 100 nucleotide segments upstream of RNAP positions, which is the length of run-on extension in GRO-seq [44]. GRO-seq tag density of each position was determined by counting the number of GRO-seq tags covering the position.

### Alignment of sequencing reads and calculation of tag density

The libraries were submitted for sequencing on the Illumina platform on one or more lanes and data has been submitted to Array Express with the following accession numbers: E-MTAB-742 for GRO-seq data, E-MTAB-743 for mRNA-seq data and E-MTAB-744 for ChIP-seq data.

All alignments were performed using the Bowtie algorithm version 0.12.5 *[45]* against hg18 assembly of human genome with a maximum of 2 mismatches per sequence allowed and only uniquely mapped reads retained. Read length and numbers and fraction of uniquely aligned reads of each channel were summarized in Table S1. Specifically for mRNA-seq, reads that failed to align to hg18 were then aligned to splice junction libraries built using Ensembl gene annotation by ERANGE version 3.2 [26]. Reads that uniquely aligned to hg18 (and splice junction libraries) in different lanes of each library were pooled to generate tag density data. To calculate tag density, aligned reads, except the ones from mRNA-seq, were extended to a fragment length of 200 nucleotides in the 5'-to-3' direction. The tag density of a genomic position was calculated as the number of tags that cover the position. For GRO-seq and mRNA-seq, tag densities of the positive strand and negative strand were calculated separately, which only counted tags aligned to the corresponding strand. In contrast, tags aligned to the two strands were pooled to calculate tag density for ChIP-seq.

### Sample Selection for Data Analysis

Total number of internal exons in Ensembl annotation was 524,514, where identical exons presenting in different isoforms were counted multiple times. Because GRO-seq tags were strongly enriched within about 1,000 bp downstream regions of 5’ ends and 10,000 bp downstream regions of 3’ ends of genes [44], we removed exons overlapping with these regions on the same strand from the Ensembl data set of internal exons. To avoid the effect of GRO-seq tag enrichment in antisense direction relative to the direction of gene transcription [44], we also excluded exons overlapping with 2,000 bp-upstream regions of 5’ ends of genes on the opposite strand. The number of unique exons that met these two criteria was 146,343. For each cell line analyzed, we further selected exons by the following two criteria. Firstly, exonic mRNA-seq tag density should be greater than 1 or equal 0. We define exons with mRNA-seq tag density greater than 1 as included exons, and equal to 0 as excluded exons. Exons with mRNA-seq tag density within (0,1] were eliminated. Secondly, for each exon, mRNA-seq tag density of at least one exon in the corresponding transcript isoforms should be greater than 1, i.e. at least one exon was included. This criterion was to ensure that the exon was in at least one expressed transcript isoform. Number of exons meeting these two additional criteria was 86,091 in IMR90, 94,638 in MCF10A, and 103,600 in ESC H1. The resulting samples of exons were used in the analysis of patterns of sequencing tag densities and de novo motif searching. In pattern score calculation, if not both exonic or intronic tag density equal 0, 0 was replaced by 0.0001, while if exonic and intronic tag density both equal 0, the exon was eliminated from relative analysis ‐‐ relationship between GRO-seq pattern score, H3K36me3 pattern score, RNAP pattern score and RNA level, and comparison of number of known splicing factor motifs among cell lines

In the random forest models (described in detail below), we used sample of IMR90 exons further selected by the criterion of exon length being greater than 200 bp. The reason for the 200bp cutoff in this analysis was that we observed strong linear correlation between exon length and H3K36me3 and weak linear correlation between exon length and H4K20me1, H3K9me3, RNAP density when exon length is smaller than about 200 bp. This correlation seems to be a result of the low resolution of ChIP-seq and the length of nucleosome footprint being the resolution limit in these experiments. In addition, DNA methylation level exhibited similarly strong correlation with exon length, as nucleosomes may be *in vivo* substrate for the DNA methyltransferases [19]. In order to limit these artificial associations due to limited resolution and to nucleosomes being the smallest organizing unit and substrate for biomolecular processes in the genome, exons with length smaller than 200 bp were removed from the sample for random forest models, resulting in a sample size of 10,779 exons in IMR90.

### Random Forest Model of Exon Inclusion

Random forest is a widely used regression and classification method [46]. In our analysis, random forest classification models were built using ‘randomForest’ package version 4.6-2 [47] with R version 2.12.1. We have defined the response binary variable, ‘exon inclusion’, as 1 if exonic mRNA-seq tag density is greater than 1, and as 0 if exonic mRNA-seq tag density is 0. The full model contains 13 continuous predictor variables shown in Table 1. Variables with ‘tag density’ in their names are exonic density of corresponding sequencing tags, ‘DNA methylation level’ is exonic density of methylcytosines reported by Lister et al. [18], variables with ‘pattern score’ in their names are pattern scores of corresponding sequencing tag density or methylcytosines density, and ‘exon length’ is the length of the corresponding exon. Because alternative splicing of internal exons is not known to be regulated by exon length, we included ‘exon length’ in the full model as a negative control to select variables with high predictive power, i.e. importance. The data set was randomly divided into 50% training and 50% test. Model was built on training set, and its performance was assessed by prediction accuracy on test set. Variable importance of the model was measured in “mean decrease in accuracy over all classes” [47]. Mean and variance of prediction accuracy and variable importance were measured by randomly resampling training and test sets 10 times. In backward variable selection starting from the full model, variable with the lowest importance (mean of 10 repeated sampling) was removed in each iteration. Backward variable selection was performed until ‘exon length’ was removed. Re-fitting the model with the remaining variables resulted in the reduced model.

### Searching for Known Splicing Factor Motifs

Putative motifs of know splicing factors were identified using the SFmap [32,33] with scoring function WR, window size 50 and high stringency. Sequence conservation was not considered. For both exonic and intronic motifs, only those within 100 bp upstream or downstream of splice sites were considered. Length-normalized exonic frequency, i.e. density, of each motif was calculated by dividing the number of the putative motifs in exonic region by the exonic length (≤ 200 bp). Similarly, intronic density of each motif was calculated by dividing the number of the putative motifs in intronic region by intronic length (200 bp).

### De Novo Motif Identification

Frequencies of nucleotide 6-, 7‐ and 8-mers generated by exhaustive enumeration of conformations were tested by Wilcoxon rank sum test to identify overrepresented ones within the splice site-adjacent regions of exons with cell type-specific GRO-seq peaks. Samples of exons with exonic GRO-seq tag enrichment in cell line *a* compared to cell line *b* was selected by the following criteria: the exon is in group 1 or 2 of cell line *a*, and in group 8, 9 or 10 of cell line *b*. Significance of p values was estimated by Benjamini and Hochberg false discovery rate (FDR) procedure [48]. Motif trees of significant n-mers (motifs) with FDR < 0.05 in each comparison were generated by STAMP [37], and motifs with distance values less than 0.05 were grouped into one cluster. Significant matches to known cis-regulatory motifs of individual motif clusters were identified by TOMTOM (version 4.6.1, default settings) [49] using TRANSFAC database (version 10.2) [35] containing 811 motifs and JASPAR Core database (2009 release) [36] containing 476 motifs. We report the results for IMR90 vs ESC H1 and MCF10A vs ESC H1; comparisons between IMR90 vs MCF10A did not reveal any significant difference in motif occurrence within either adjacent exonic regions or intronic regions of splice sites.

## ACKNOWLEDGMENTS

We would like to thank Joan Steitz and the members of our laboratories for critical comments on the manuscript and helpful discussions.

## SUPPORTING FIGURE LEGENDS

**Figure S1.** GRO-seq tag density patterns in MCF10A.

GRO-seq reads aligned around exon start sites (upper panel), midpoints (middle panel) and end sites (lower panel) whose positions are shown by vertical dashed lines. Red curves and blue curves correspond to exons on positive strand and negative strand, respectively. Three distinct patterns (dip, flat behavior and peak) are observed within exons in groups with low (group 1 to 5), medium (group 6) and high (group 7 to 10) exonic GRO-seq tag densities, while intronic GRO-seq tag densities are about the same (except group 1 and 10).

**Figure S2.** GRO-seq tag density patterns in IMR90.

GRO-seq reads aligned around exon start sites (upper panel), midpoints (middle panel) and end sites (lower panel) whose positions are shown by vertical dashed lines. Red curves and blue curves correspond to exons on positive strand and negative strand, respectively. Three distinct patterns (dip, flat behavior and peak) are observed within exons in groups with low (group 1 to 5), medium (group 6) and high (group 7 to 10) exonic GRO-seq tag densities, while intronic GRO-seq tag densities are about the same (except group 1 and 10).

**Figure S3.** GRO-seq tag density patterns in H1ESC.

GRO-seq reads aligned around exon start sites (upper panel), midpoints (middle panel) and end sites (lower panel) whose positions are shown by vertical dashed lines. Red curves and blue curves correspond to exons on positive strand and negative strand, respectively. Three distinct patterns (dip, flat behavior and peak) are observed within exons in groups with low (group 1 to 5), medium (group 6) and high (group 7 to 10) exonic GRO-seq tag densities, while intronic GRO-seq tag densities are about the same (except group 1 and 10).

**Figure S4.** Exons with different GRO-seq tag density patterns do not differ in GC content.

GC content around exon start sites (upper panel), midpoints (middle panel) and end sites (lower panel), whose positions are indicated by dashed vertical lines, are shown for 10 groups of exons divided by exonic GRO-seq tag density of IMR90. Specifically, GC content and T content are percentages of nitrogenous bases (G and C) in every 5 base pairs.

**Figure S5.** Exons with different GRO-seq tag density patterns do not differ T content.

T content around exon start sites (upper panel), midpoints (middle panel) and end sites (lower panel), whose positions are indicated by dashed vertical lines, are shown for 10 groups of exons divided by exonic GRO-seq tag density of IMR90. Specifically, T content are percentages of thymine (T) in every 5 base pairs.

**Figure S6.** Exons with different GRO-seq tag density patterns do not differ in mapability.

Mapability around exon start sites (upper panel), midpoints (middle panel) and end sites (lower panel), whose positions are indicated by dashed vertical lines, are shown for 10 groups of exons divided by exonic GRO-seq tag density of IMR90. Mapability is the value of Duke 35 bp Uniqueness in single base-pair resolution downloaded from UCSC Genome Browser (*18*), shown in the scale of 0 to 1.

**Figure S7.** GRO-seq pattern score distributions.

Pattern score distributions for each of the ten groups in (A) MCF10, (B) IMR90, and (C) H1ESC cell lines are shown. In each panel, a red dashed vertical line indicate where pattern score equals 1.

**Figure S8.** GRO-seq pattern score distributions.

Histogram of all GRO-seq pattern scores for all examined exons in IMR90. Frequency is noted in the Y-xis and the GRO-seq pattern score is noted in the X-axis

**Figure S9.** GRO-seq tag density pattern score correlates with mRNA level in IMR90.

Bar plot shows distributions of GRO-seq pattern scores in 10 groups divided by exonic mRNA-seq tag density. Each group of mRNA-seq tag density is represented by one bar. Exons were also classified into 10 equal-sized groups of continuous value ranges of GRO-seq pattern score, as indicated by the legend on the right of the plot. The proportions of exons within different GRO-seq pattern score value ranges show a shift among 10 groups of mRNA-seq tag density.

**Figure S10.** GRO-seq tag density pattern score correlates with mRNA level in H1ESC.

Bar plot shows distributions of GRO-seq pattern scores in 10 groups divided by exonic mRNA-seq tag density. Each group of mRNA-seq tag density is represented by one bar. Exons were also classified into 10 equal-sized groups of continuous value ranges of GRO-seq pattern score, as indicated by the legend on the right of the plot. The proportions of exons within different GRO-seq pattern score value ranges show a shift among 10 groups of mRNA-seq tag density.

**Figure S11.** GRO-seq pattern scores distinguish excluded from included exons.

Distribution of GRO-seq pattern score over excluded exons (left panel) is different from which of moderately expressed (middle panel) and highly expressed (right panel) exons in MCF10A with respect to slope of distribution function. In each panel, a red dashed vertical line indicate where pattern score equals 1.

**Figure S12.** RNAP pattern scores are not related with mRNA level in MCF10A.

(A) Bar plot shows distributions of RNAP-seq pattern scores in 10 groups divided by exonic mRNA-seq tag density. Each group of mRNA-seq tag density is represented by one bar. Exons were also classified into 10 equal-sized groups of continuous value ranges of RNAP-seq pattern score, as indicated by the legend on the right of the plot. The proportions of exons within different RNAP-seq pattern score value ranges show a shift among 10 groups of mRNA-seq tag density.

**Figure S13.** RNAP pattern scores do not distinguish excluded and included exons.

Histograms of distributions of RNAP pattern score over excluded, moderately expressed and highly expressed exons. In each panel, a red dashed vertical line indicate where pattern score equals 1.

**Figure S14.** Distribution of GRO-seq pattern scores at included and excluded exons of the same gene.

(A) Binned density distribution of GRO-seq pattern scores at included exons (Y-axis) and excluded exons (X-axis) of the same genes. Each hexagon is shaded according to the gray scale count legend on the right. The included exons have higher GRO-seq pattern score than excluded exons. (B) Distributions of the difference of GRO-seq tag densities found at the included and excluded exons of the same gene. The rightward skew of the distribution indicates that included exons exhibit higher GRO-seq tag density than the excluded exons of the same gene.

**Figure S15.** H3K36me3 tag density pattern does not explain GRO-seq tag density pattern.

Distributions of H3K36me3 pattern score over 10 groups of exons divided by exonic GRO-seq tag density do not show strong difference between groups, in comparison with the shift of the GRO-seq pattern score (Figure S2B).

**Figure S16.** H3K36me3 pattern scores are not related with mRNA level in MCF10A.

Bar plot shows distributions of H3K36me3 ChIP-seq pattern scores in 10 groups divided by exonic mRNA-seq tag density. Each group of mRNA-seq tag density is represented by one bar. Exons were also classified into 10 equal-sized groups of continuous value ranges of H3K36me3 ChIP-seq pattern score, as indicated by the legend on the right of the plot. The proportions of exons within different H3K36me3 ChIP-seq pattern score value ranges show a shift among 10 groups of mRNA-seq tag density.

**Figure S17.** H3K36me3 pattern scores do not distinguish excluded from included exons.

Histograms of distributions of H3K36me3 pattern score over excluded, medium expressed and highly expressed exons. In each panel, a red dashed vertical line indicate where pattern score equals 1.

**Figure S18.** Sample selection for splicing factor motif analysis.

In the comparison “a vs b” (Fig. 4), sample a contains exons with high exonic GRO-seq tag densities in cell type a and low exonic GRO-seq tag densities in cell type b, sample b contains exons with low exonic GRO-seq tag densities in cell type a and high exonic GRO-seq tag densities in cell type b, while other exons are eliminated (see Methods and Materials).

**Figure S19**: Differential enrichments of putative known splicing factor motifs in exons with cell-type-specific GRO-seq patterns.

P values of one-sided Wilcoxon rank sum tests for length-normalized frequencies, i.e. densities, of exonic (A) or intronic (B) splicing factor motifs located no further than 100 bp from splicing sites. Cell types being compared are IMR90, MCF10A and ESC H1. For each of the three panels, the two cell types being compared are indicated on the top in the form of “a vs b”, with sample sizes denoted by n. Wilcoxon rank sum tests are performed between exonic and intronic motif densities of exons with exonic GRO-seq pattern scores > 2 in cell type “a” but ≤ 0.5 in cell type “b” and those with exonic GRO-seq pattern scores ≤ 0.5 in cell type “a” but > 2 in cell type “b”. The p value of each test is shown in the palette. Significant p values were further defined by Benjamini and Hochberg FDR procedure(*19*) at the 0.05 level and indicated by colored background: yellow means motifs are enriched in cell type “a”, blue means motifs are enriched in cell type “b”, and grey means no significant enrichment.

